# Muscle-derived miR-200a-3p through light-intensity exercise may contribute to improve memory dysfunction in type 2 diabetic mice

**DOI:** 10.1101/2024.05.24.595713

**Authors:** Takeru Shima, Hayate Onishi, Chiho Terashima

## Abstract

**Background:** Memory dysfunction associated with type 2 diabetes mellitus (T2DM) poses a considerable threat to overall well-being. Engaging in light-intensity exercise has been shown to exert favorable effects on hippocampal function and molecular profiles, including *Mct2* mRNA and miR-200a-3p. Nonetheless, a comprehensive understanding of the mechanisms underlying the positive impact of light-intensity exercise remains elusive. Here, we assessed the influence of exosomal miR-200a-3p secretion from gastrocnemius muscles in T2DM mice undergoing light-intensity exercise intervention, focusing on its potential to ameliorate memory dysfunction.

**Basic procedures:** We initially assessed the effects of light-intensity exercise (7.0 m/min for healthy mice, 5.0 m/min for ob/ob mice, 30 min/day, five days/week, over a four-week period) on memory function, hippocampal mRNA associated with memory function, and the secretion of exosomal miR-200a-3p from their gastrocnemius muscle. Subsequently, the impact of a daily intraperitoneal injection of the miR-200a-3p mimic over a four-week duration was investigated, focusing on its influence on hippocampal dysregulation in ob/ob mice.

**Main findings:** The light-intensity exercise intervention increased gastrocnemius muscle-derived and plasma exosomal miR-200a-3p levels in ob/ob mice, concomitant with improved memory dysfunction. Intriguingly, the daily intraperitoneal injection of mmu-miR-200a-3p mimic also demonstrated an ameliorative effect on memory function in ob/ob mice. Notably, both the exercise intervention and miR-200a-3p mimic treatment induced downregulation in hippocampal *Keap1* mRNA and upregulation in mRNA of *Hsp90aa1* and *Mct2* in ob/ob mice.

**Principal conclusions:** The current results imply that the augmentation of exosomal miR-200a-3p derived from the gastrocnemius muscle contributes to the amelioration of memory dysfunction in T2DM undergoing light-intensity exercise. Additionally, it is proposed that miR-200a-3p emulates the effects of light-intensity exercise, suggesting a potential therapeutic pathway for addressing hippocampal complications in the context of T2DM.

## 1. Introduction

In light of the favorable impact that physical exercise exerts on both peripheral organs and the brain, it has emerged as a widely adopted therapeutic strategy for addressing metabolic syndrome and mental health disorders [1]. Within the context of type 2 diabetes (T2DM), exercise therapy primarily focuses on mitigating insulin resistance and enhancing glycemic control [2,3]. T2DM triggers various complications not only in peripheral organs but also in the hippocampus, leading to memory dysfunction [4–6]. It is anticipated that exercise can serve as a therapeutic intervention for memory dysfunction in T2DM [7]. Therefore, elucidating the mechanisms underlying the effects of exercise on the hippocampus in T2DM holds the potential to develop a therapeutic approach targeting the hippocampus in T2DM, a crucial endeavor for the preservation of human well-being.

Downregulation of lactate transport through monocarboxylate transporter 2 (MCT2) into neurons is assumed to be a potential etiological mechanism underlying T2DM-induced memory dysfunction [8–11]. Lactate is a crucial energy substrate for neurons, derived from blood circulation and astrocytes through glycolysis and glycogenolysis pathways [12–15]. Within astrocytes, glycogen functions as a principal reservoir for lactate. The release of glycogen-derived lactate from astrocytes into the extracellular fluid is facilitated by transporters MCT1 and MCT4, subsequently taken up by neurons through MCT2 [15]. Neurons utilize lactate both as an energy substrate for their activity and as a neuromodulator, contributing to neuronal plasticity [16–18]. Previous studies have demonstrated that hindering lactate supply and downregulating MCT2 expression and function within the hippocampus result in impairments in learning and memory [16,19–22]. Significantly, rodents with T2DM exhibit reduced expression levels of MCT2 in their hippocampi, without notable differences in MCT1 or MCT4 expressions compared to control rodents [8–11]. Research has shown that a light-intensity exercise intervention improves hippocampal memory dysfunction and enhances MCT2 expression in rodent models of T2DM [11,23]. Consequently, the compromised transport of lactate through MCT2 likely plays a pivotal role in the hippocampal complications associated with memory dysfunction in T2DM, shedding light on the potential therapeutic effects of exercise.

Several studies have proposed that the release of exosomes constitutes a fundamental mechanism underpinning the effects of exercise. These exosomes, characterized as small extracellular vesicles within the range of 40 to 160 nm, are released by diverse organs and encapsulate microRNAs (miRNAs). These miRNAs, short non-coding RNAs measuring approximately 21-25 nucleotides, engage in binding with complementary segments of messenger RNAs (mRNAs), thereby initiating mRNA degradation or hindering translation [24]. Notably, emerging evidence suggests that exercise-induced exosomal miRNAs, originating from muscles, contribute to enhanced biological and physiological functions in peripheral organs and the brain [25–28]. However, it remains unclear whether exosomal miRNAs released during light-intensity exercise are associated with the improvement of hippocampal function in the context of T2DM.

A prior investigation has documented that light-intensity exercise intervention increases miR-200a-3p levels, accompanied by the upregulation of *Mct2* mRNA levels in the hippocampus [11]. MiR-200a-3p is notable for its role in cell growth and its inhibitory effect on mRNA levels in Kelch ECH-associated protein 1 (*Keap1*) and phosphatase and tensin homolog (*Pten*) [29,30]. KEAP1 functions as a negative regulator for nuclear factor erythroid 2 p45-related factor-2 (NRF2) [31] and heat shock protein 90 (HSP90) [32]. In the context of T2DM, there is observed overexpression of KEAP1 in various organs, including the hippocampus, which is hypothesized to be a potential contributor to complications associated with T2DM [33–35]. Notably, it has been reported that high-intensity interval exercise intervention improves the overexpression of KEAP1 in the T2DM hippocampus [35]. Furthermore, PTEN is implicated in T2DM pathogenesis [36] and has an impact on the expressions of brain-derived neurotrophic factor (BDNF) [37]. Previous studies have proposed potential associations between profiles of HSPs and the expressions of MCTs [38,39], while BDNF is known to modulate MCT2 expressions [40,41]. Thus, the restoration of miR-200a-3p levels in the hippocampus may contribute to the amelioration of memory dysfunction and the enhancement of hippocampal MCT2 expressions in T2DM through the modulation of mRNA levels in *Keap1* and *Pten*. Furthermore, a previous study has indicated that 4 weeks of light-intensity exercise did not significantly alter circulating miR-200a-3p levels in healthy mice [42]; however, the impact of such exercise intervention on circulating miR-200a-3p levels in T2DM mice remains uncertain.

Here, we initially evaluated the impact of light-intensity exercise intervention on the release of exosomal miR-200a-3p from gastrocnemius muscles in T2DM mice, simultaneously addressing improvements in memory dysfunction and alterations in hippocampal mRNA expressions. Subsequently, we conducted a detailed investigation into the effects of daily intraperitoneal injections of mmu-miR-200a-3p mimic on both hippocampal memory function and mRNA expressions in T2DM mice.

## 2. Methods and materials

### 2.1. Animals

Male C57BL/6 mice and ob/ob mice (a T2DM mouse model) at eight weeks of age were obtained from SLC Inc. (Japan) and housed in a temperature-controlled room (21-23□) with a 12-hour light/dark cycle (lights on from 7 AM to 7 PM). They were provided ad libitum access to a standard pellet diet (Rodent Diet CE-2, CLEA Japan Inc., Japan) and water. The experimental procedures were pre-approved (approval No. 22-012) and conducted in accordance with the guidelines established by the Gunma University Animal Care and Experimentation Committee.

### 2.2. Exercise training

Following a week of acclimatization, both C57BL/6 mice and ob/ob mice at 9 weeks of age were divided into exercise and non-exercise (sedentary) groups, ensuring a match in body weight between the groups. The groups were as follows: sedentary C57BL/6 (n = 10), exercised C57BL/6 (n = 9), sedentary ob/ob (n = 9), and exercised ob/ob (n = 9). Mice in the exercise group underwent a running habituation phase on a forced exercise wheel bed for 30 min/day, five days/week (a total of five sessions over six days) at speeds ranging from 3.0 to 7.0 m/min for C57BL/6 mice and from 3.0 to 5.0 m/min for ob/ob mice, spanning one week. Subsequently, they engaged in light-intensity exercise sessions (C57BL/6 mice, 7.0 m/min; ob/ob mice, 5.0 m/min) on the same equipment for 30 min/day, five days/week over three weeks [11,43,44]. The exercise intensity for each strain was established based on their ventilatory threshold [44]. All sessions were conducted during the light period (from 7 AM to 9 AM). Memory performance tests were conducted for both exercise and sedentary groups over a period of five days during the final week of the exercise regimen.

### 2.3. Intraperitoneal injection of mmu-miR-200a-3p mimic

After one week of acclimatization, the mice at 9 weeks of age were divided into miR-200a-3p mimic-treated and miRNA mimic negative control-treated groups, ensuring that body weights were matched between the groups. Mice in miR-200a-3p mimic-treated group and miRNA mimic negative control-treated group received daily intraperitoneal injections of mmu-miR-200a-3p mimic (Ajinomoto Bio-Pharma, Japan) and miRNA mimic negative control (mimic NC; SMC-2003, Bioneer, Korea) for 4 weeks, respectively. A 20 nmol/l solution of each drug was prepared by diluting it in 0.9% saline with 0.02% TE buffer. Mice received an injection of 10 μl/g body weight of either the mmu-miR-200a-3p mimic or the mimic NC solution during the light period (0.2 nmol/kg body weight, once a day from 8 AM to 9 AM). The designated groups were as follows: C57BL/6 mimic NC (n = 8), C57BL/6 miR-200a-3p mimic (n = 8), ob/ob mimic NC (n = 8), and ob/ob miR-200a-3p mimic (n = 8). These mice did not undergo exercise training. Memory performance tests were conducted for all mice over a period of five days during the final week of the treatment.

### 2.4. Memory performance test

The Morris water maze test was conducted in a circular pool (100 cm in diameter and 30 cm in depth) with an invisible platform (10 cm in diameter) positioned in the center of one quadrant. The experimental room had several extra-maze cues. All four start-points were utilized during the learning sessions in different sequences. Mice were given 60 seconds to explore and locate the platform. In cases where mice failed to find the platform within 60 seconds, they were manually guided to it. Upon reaching the platform, the mice remained there for 10 seconds. Throughout the learning sessions, escape latency (sec), swim length (cm), and speed (cm/sec) were recorded using a video tracking system (O’hara & Co., Ltd., Japan). One day after the final learning session, the platform was removed from the pool, and mice underwent a probe trial for 60 seconds to search for it within the pool. The time spent in the quadrants where the platform had been located during the learning sessions was measured using the same tracking system.

### 2.5. Tissue preparation

Two days following the probe trial, mice were anesthetized using isoflurane (Dainippon Sumitomo Pharma Co., Japan), and the blood samples were obtained from cardiac puncture. Subsequently, the hippocampus was collected and preserved in RNAlater™ Stabilization Solution (Invitrogen™, USA). The hippocampal tissues were stored at -20□ for subsequent biochemical analysis. Furthermore, gastrocnemius muscle and plasma were collected and used to extract exosomes.

### 2.6. Extraction of RNAs and exosomal miRNAs

Total RNAs and miRNAs were extracted from the hippocampal tissue using the RNeasy Mini Kit and the miRNeasy Micro Kit (Qiagen Inc., USA), respectively. Gastrocnemius muscle tissues obtained from mice were promptly incubated in DMEM (Gibco-Thermo Fisher Scientific Inc., USA) supplemented with 10% of exosome-depleted FBS (EXO-FBSHI-50A-1; SBI LLC., USA) and 1% of Pen-Strep-Glutamine (Gibco-Thermo Fisher Scientific Inc., USA) for 24 hours in a humidified incubator at 37°C and 5% CO_2_. The medium was then collected and filtered through 22 μm filters. Gastrocnemius muscle-derived exosomal miRNAs in the medium were extracted using exoRNeasy Midi Kit (Qiagen Inc., USA). Exosomal miRNAs in plasma samples were also extracted using exoRNeasy Midi Kit (Qiagen Inc., USA). As per the protocol provided from Qiagen Inc (USA), 150 μl of plasma in each mouse was utilized to extract exosomal miRNAs.

### 2.7. Real-time PCR

After extracting RNAs from the hippocampus, DNase I treatment was applied, and RNA quantification was carried out using the Qubit 4.0 (Invitrogen™, USA). For the detection of mRNA levels in the hippocampus, 1000 ng of RNA underwent reverse transcription to cDNA using the GeneAce cDNA Synthesis Kit (Nippon Gene, Japan). Subsequently, the mRNA levels of target genes were assessed using 5.0 ng of cDNA, primers for each target gene, and the PowerTrack^™^□ SYBR™□ Green Master Mix in the StepOne Plus Real-Time PCR 96-well system (Thermo Fisher Scientific Inc., USA). The primer sequences (forward and reverse) used in the current study are provided in Table S1. The relative levels of each mRNA were calculated utilizing the ΔΔCT method and normalized by β-actin mRNA levels.

For the detection of miRNA levels in the hippocampus, as well as in gastrocnemius muscle- and plasma-derived exosomes, 10 ng of miRNA underwent reverse transcription to cDNA using the Taqman™□ MicroRNA Reverse Transcription Kit and the Taqman™□ MicroRNA Assay (miR-200a-3p: 000502, and U6 snRNA: 001973; Thermo Fisher Scientific Inc., USA). Subsequently, miR-200a-3p and U6 levels were quantified using 0.67 ng of cDNA, the Taqman™□ MicroRNA Assay, and the Taqman™□ Fast Advanced Master Mix in the StepOne Plus (Thermo Fisher Scientific Inc., USA). The relative levels of miR-200a-3p were calculated by ΔΔCT method and normalized by U6 snRNA levels.

### 2.8. Statistical analysis

The data are presented as mean ± standard error (SEM) and were analyzed using Prism version 10.1.1 (MDF, Japan). Group comparisons were conducted utilizing repeated three-way ANOVA (factor 1: day in the Morris water maze test; factor 2: diabetes [C57BL/6 mice vs. ob/ob mice]; factor 3: exercise [sedentary vs. exercised] or miR-200a-3p [mimic NC vs. miR-200a-3p mimic]) or two-way ANOVA (factor 1: diabetes [C57BL/6 mice vs. ob/ob mice]; factor 2: exercise [sedentary vs. exercised] or miR-200a-3p [mimic NC vs. miR-200a-3p mimic]) with Tukey’s post hoc tests. We used repeated three-way ANOVA solely for analyzing the results of escape latency, swim length, and speed during the learning sessions of the Morris water maze test. Correlations were analyzed by Pearson correlation. Statistical significance was set at *p* < 0.05.

## 3. Results

### 3.1. The effects of light-intensity exercise on physiological and biochemical variables

Both sedentary and exercised ob/ob mice exhibited significantly higher body weight and fat-to-body weight ratio compared to both sedentary and exercised C57BL/6 mice (Table S2; body weight, effects of diabetes: *F*_(1, 33)_ = 1353.6, *p* < 0.0001, effects of exercise: *F*_(1, 33)_ = 0.99, *p* = 0.3280, interaction: *F*_(1, 33)_ = 0.78, *p* = 0.3831; fat-to-body weight ratio, effects of diabetes: *F*_(1, 33)_ = 2046.9, *p* < 0.0001, effects of exercise: *F*_(1, 33)_ = 3.05, *p* = 0.0899, interaction: *F*_(1, 33)_ = 1.25, *p* = 0.2711). Sedentary ob/ob mice exclusively demonstrated elevated blood glucose levels in comparison to both sedentary and exercised C57BL/6 mice, whereas exercised ob/ob mice did not exhibit this distinction (effects of diabetes: *F*_(1, 33)_ = 15.33, *p* = 0.0004, effects of exercise: *F*_(1, 33)_ = 2.73, *p* = 0.1081, interaction: *F*_(1, 33)_ = 0.88, *p* = 0.3548). Although HbA_1C_ levels were markedly higher in both groups of ob/ob mice compared to sedentary and exercised C57BL/6 mice, those in exercised ob/ob mice were significantly lower than those in sedentary ob/ob mice (effects of diabetes: *F*_(1, 33)_ = 344.4, *p* < 0.0001, effects of exercise: *F*_(1, 33)_ = 12.55, *p* = 0.0012, interaction: *F*_(1, 33)_ = 11.50, *p* = 0.0018).

### 3.2. The effects of light-intensity exercise on memory function

Sedentary ob/ob mice exhibited significantly prolonged escape latency compared to both sedentary and exercised C57BL/6 mice, whereas exercised ob/ob mice did not exhibit this difference on 2^nd^ and 4^th^ day of learning session (Fig. 1A; effects of day: *F*_(3, 432)_ = 15.57, *p* < 0.0001, effects of diabetes: *F*_(1, 144)_ = 62.86, *p* < 0.0001, effects of exercise: *F*_(1, 144)_ = 6.59, *p* = 0.0113, interaction [day × diabetes]: *F*_(3, 432)_ = 3.68, *p* = 0.0122, interaction [day × exercise]: *F*_(3, 432)_ = 2.03, *p* = 0.1088, interaction [diabetes × exercise]: *F*_(1, 144)_ = 0.11, *p* = 0.7370, interaction [day × diabetes × exercise]: *F*_(3, 432)_ = 1.09, *p* = 0.3524). Swim distance and swimming speed were influenced by time and T2DM but not by exercise (Fig. 1B and C; swim distance, effects of day: *F*_(3, 432)_ = 22.76, *p* < 0.0001, effects of diabetes: *F*_(1, 144)_ = 8.51, *p* = 0.0041, effects of exercise: *F*_(1, 144)_ = 2.42, *p* = 0.1219, interaction [day × diabetes]: *F*_(3, 432)_ = 3.17, *p* = 0.0244, interaction [day × exercise]: *F*_(3, 432)_ = 0.46, *p* = 0.7132, interaction [diabetes × exercise]: *F*_(1, 144)_ = 1.21, *p* = 0.2734, interaction [day × diabetes × exercise]: *F*_(3, 432)_ = 0.84, *p* = 0.4704; swimming speed, effects of day: *F*_(3, 432)_ = 9.28, *p* < 0.0001, effects of diabetes: *F*_(1, 144)_ = 116.9, *p* < 0.0001, effects of exercise: *F*_(1, 144)_ = 0.35, *p* = 0.5579, interaction [day × diabetes]: *F*_(3, 432)_ = 1.16, *p* = 0.3239, interaction [day × exercise]: *F*_(3, 432)_ = 1.32, *p* = 0.2685, interaction [diabetes × exercise]: *F*_(1, 144)_ = 2.55, *p* = 0.1127, interaction [day × diabetes × exercise]: *F*_(3, 432)_ = 1.87, *p* = 0.1342). During the probe test, the time spent in the platform area by exercised ob/ob mice was significantly greater than that observed in sedentary C57BL/6 mice and ob/ob mice, respectively (Fig. 1D; effects of diabetes: *F*_(1, 33)_ = 0.001, *p* = 0.9731, effects of exercise: *F*_(1, 33)_ = 17.49, *p* = 0.0002, interaction: *F*_(1, 33)_ = 7.17, *p* = 0.0115). In contrast, sedentary ob/ob mice exhibited significantly shorter times spent in the platform area compared to exercised C57BL/6 mice (Fig. 1D).

**Fig. 1.**
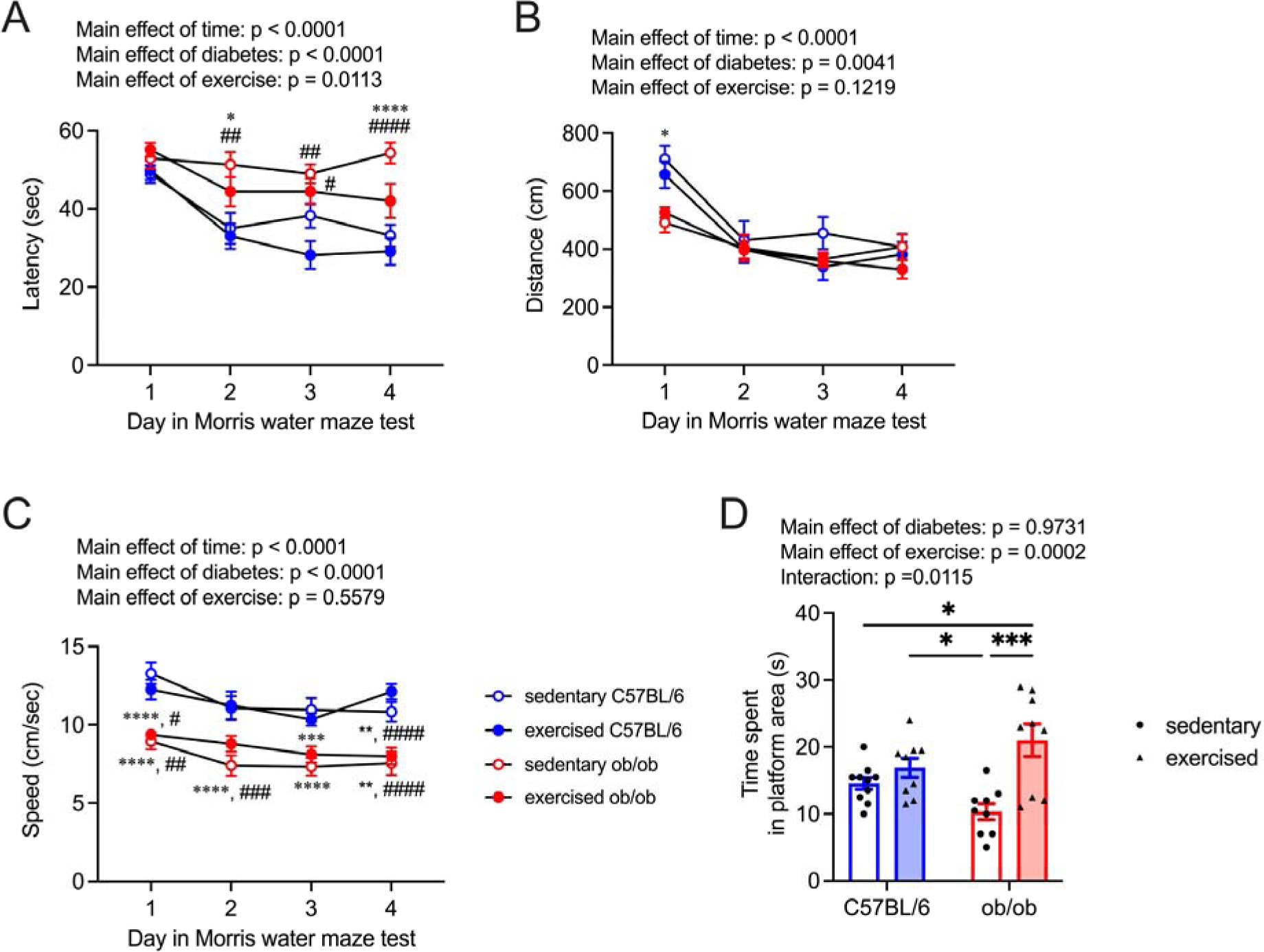
Effect of light-intensity exercise on memory function. Escape latency (A), swim length (B), and speed (C) during the learning session in mice (mean ± SEM). Blue circles: C57BL/6 mice, red circles: ob/ob mice. \*\**p* < 0.01 vs sedentary C57BL/6, ^#^*p* < 0.05, ^##^*p* < 0.01 vs exercised C57BL/6. (D) Effect of exercise on the probe trial, showing the crossing times where the platform had been placed. Blue bars: C57BL/6, red bars: ob/ob, circles: sedentary groups, and triangles: exercised groups. \**p* < 0.05, \*\*\**p* < 0.001. Data are expressed as mean ± SEM, n = 10 mice for sedentary C57BL/6, and n = 9 mice for exercised C57BL/6, sedentary ob/ob, and exercised ob/ob.

### 3.3. The effects of light-intensity exercise on mRNA levels related to lactate transport in the hippocampus

The sedentary ob/ob mice exhibited notably reduced hippocampal *Mct2* mRNA levels compared to both sedentary and exercised C57BL/6 mice; however, this difference was not observed in exercised ob/ob mice (Fig. 2B; effects of diabetes: *F*_(1, 33)_ = 9.69, *p* = 0.0038, effects of exercise: *F*_(1, 33)_ = 0.88, *p* = 0.3553, interaction: *F*_(1, 33)_ = 2.32, *p* = 0.1370). The mRNA levels of *Mct1* and *Mct4* in the hippocampus remained unaltered regardless of T2DM or exercise (Fig. 2A and C; *Mct1*, effects of diabetes: *F*_(1, 33)_ = 2.95, *p* = 0.0952, effects of exercise: *F*_(1, 33)_ = 0.69, *p* = 0.4125, interaction: *F*_(1, 33)_ = 1.71, *p* = 0.2003; *Mct4*, effects of diabetes: *F*_(1, 33)_ = 0.08, *p* = 0.7762, effects of exercise: *F*_(1, 33)_ = 0.10, *p* = 0.7561, interaction: *F*_(1, 33)_ = 0.03, *p* = 0.8558). In contrast, hippocampal *Hcar1* mRNA levels were significantly elevated in mice with T2DM compared to control mice (Fig. 2D; effects of diabetes: *F*_(1, 33)_ = 7.93, *p* = 0.0081, effects of exercise: *F*_(1, 33)_ = 0.60, *p* = 0.4454, interaction: *F*_(1, 33)_ = 1.03, *p* = 0.3177). Both sedentary and exercised ob/ob mice showed significantly diminished *Bdnf* mRNA levels in the hippocampus compared to both sedentary and exercised C57BL/6 mice (Fig. 2E; effects of diabetes: *F*_(1, 33)_ = 31.11, *p* < 0.0001, effects of exercise: *F*_(1, 33)_ = 0.001, *p* = 0.9776, interaction: *F*_(1, 33)_ = 0.74, *p* = 0.3954). Although hippocampal tropomyosin□related kinase B (*Trkb*) mRNA levels remained unaffected by T2DM or exercise (Fig. 2F; effects of diabetes: *F*_(1, 33)_ = 2.38, *p* = 0.1323, effects of exercise: *F*_(1, 33)_ = 0.12, *p* = 0.7324, interaction: *F*_(1, 33)_ = 0.59, *p* = 0.4478), mRNA levels of cAMP response element binding protein (*Creb1*) in the hippocampus were lower in T2DM mice compared to control mice (Fig. 2G; effects of diabetes: *F*_(1, 33)_ = 5.33, *p* = 0.0274, effects of exercise: *F*_(1, 33)_ = 2.23, *p* = 0.1450, interaction: *F*_(1, 33)_ = 0.18, *p* = 0.6713).

**Fig. 2.**
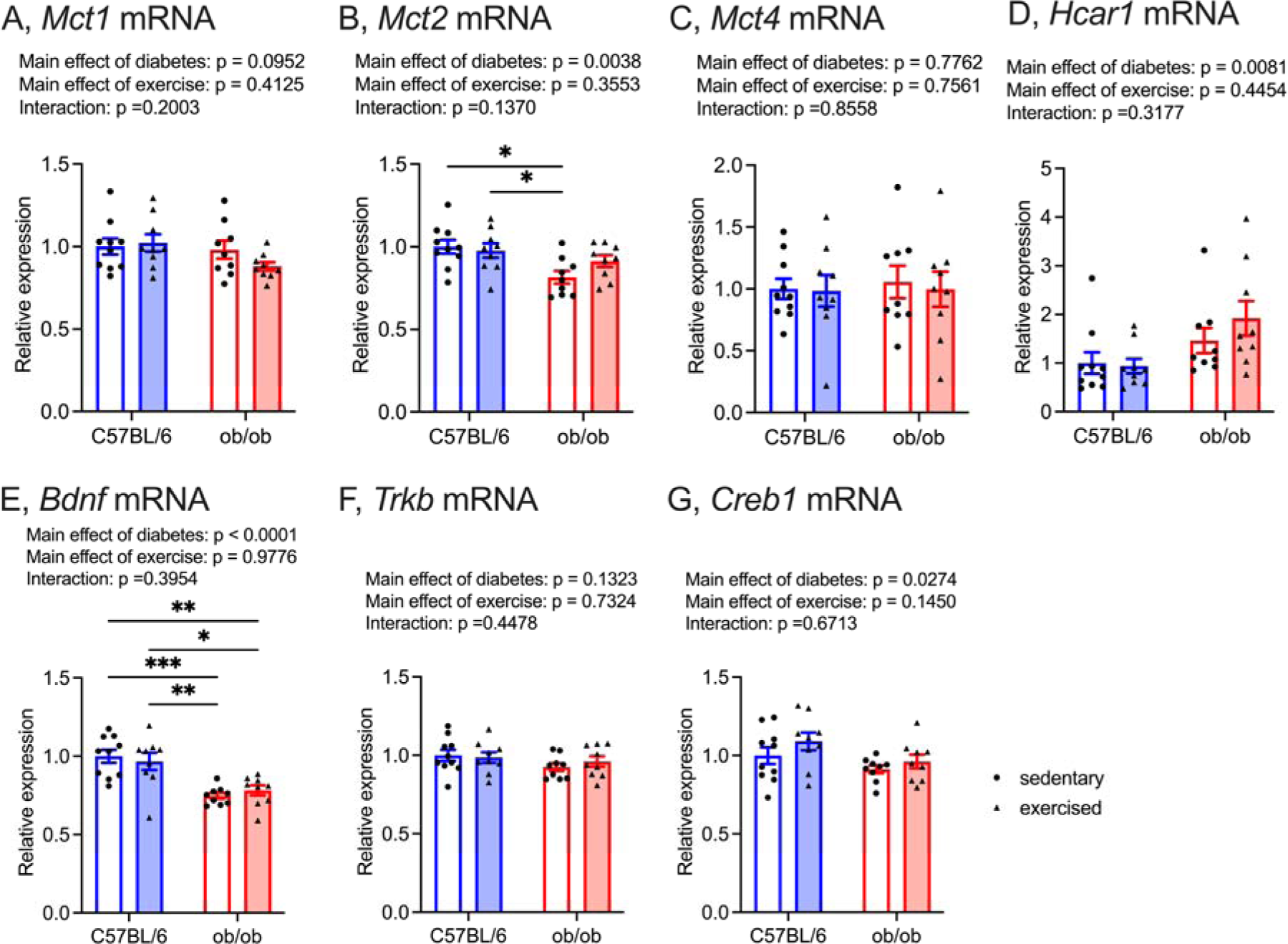
Effect of light-intensity exercise on mRNA levels of *Mct1* (A), *Mct2* (B), *Mct4* (C), *Hcar1* (D), *Bdnf* (E), *Trkb* (F), *Creb1* (G) in the hippocampus. Sedentary C57BL/6 group was normalized as 100%. Data are expressed as mean ± SEM, n = 10 mice for sedentary C57BL/6, and n = 9 mice for exercised C57BL/6, sedentary ob/ob, and exercised ob/ob. \**p* < 0.05, \*\**p* < 0.01, \*\*\**p* < 0.001.

### 3.4. The effects of light-intensity exercise on exosomal and hippocampal miR-200a-3p and its-related mRNA levels in the hippocampus

Although the downregulation of gastrocnemius muscle-derived exosomal miR-200a-3p levels in the presence of T2DM, light-intensity exercise increased these levels (Fig. 3A; effects of diabetes: *F*_(1, 33)_ = 8.47, *p* = 0.0064, effects of exercise: *F*_(1, 33)_ = 6.80, *p* = 0.0136, interaction: *F*_(1, 33)_ = 0.0001, *p* = 0.9934). Specifically, sedentary ob/ob mice exhibited significantly lower gastrocnemius muscle-derived exosomal miR-200a-3p levels compared to exercised C57BL/6 mice (Fig. 3A). Light-intensity exercise significantly enhanced exosomal miR-200a-3p levels in the plasma of ob/ob mice (Fig. 3B; effects of diabetes: *F*_(1, 33)_ = 0.97, *p* = 0.3327, effects of exercise: *F*_(1, 33)_ = 2.02, *p* = 0.1643, interaction: *F*_(1, 33)_ = 3.67, *p* = 0.0640). Hippocampal miR-200a-3p levels showed a tendency to decrease in sedentary ob/ob mice compared to sedentary C57BL/6 mice, but this tendency was not observed in exercised ob/ob mice (Fig. 3C; effects of diabetes: *F*_(1, 33)_ = 3.59, *p* = 0.0670, effects of exercise: *F*_(1, 33)_ = 0.21, *p* = 0.6529, interaction: *F*_(1, 33)_ = 2.93, *p* = 0.0963). A positive association between gastrocnemius muscle-derived exosomal miR-200a-3p levels and plasma exosomal miR-200a-3p levels was evident in ob/ob mice, but not in C57BL/6 mice (Fig. 3D). Additionally, a weak, albeit nonsignificant, correlation was observed between exosomal miR-200a-3p levels in plasma and hippocampal miR-200a-3p levels in ob/ob mice, but not in C57BL/6 mice (Fig. 3E).

**Fig. 3.**
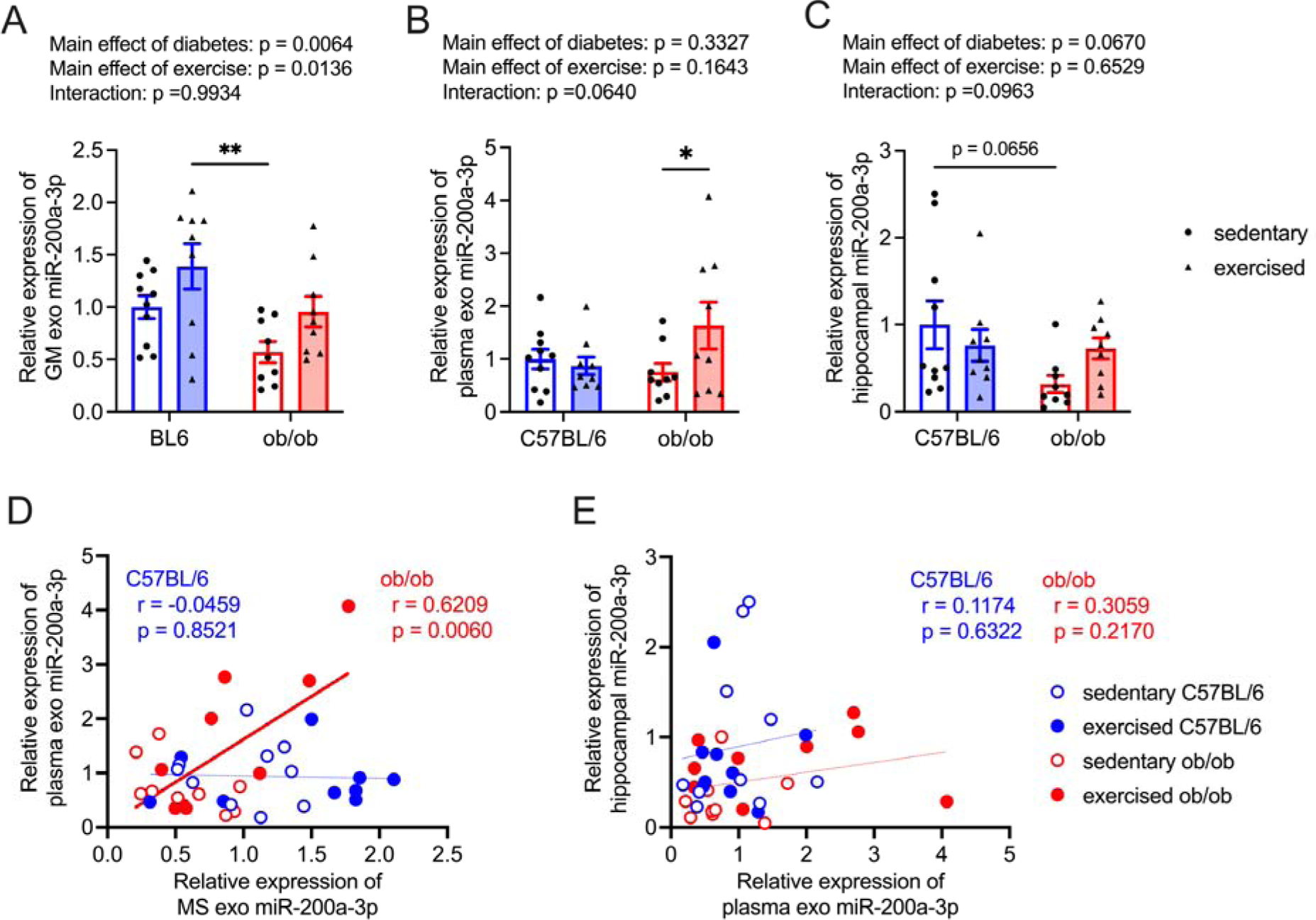
Effect of light intensity exercise on gastrocnemius muscle-derived exosomal miR-200a-3p (A), exosomal miR-200a-3p in plasma (B), and miR-200a-3p in hippocampus (C). Sedentary C57BL/6 group was normalized as 100%. Data are expressed as mean ± SEM, n = 10 mice for sedentary C57BL/6, and n = 9 mice for exercised C57BL/6, sedentary ob/ob, and exercised ob/ob. \**p* < 0.05, \*\**p* < 0.01. (D) The correlations between gastrocnemius muscle-derived exosomal miR-200a-3p and exosomal miR-200a-3p in plasma. The red line in the scatter diagram indicates significant correlation in ob/ob mice. (E) The correlations between exosomal miR-200a-3p in plasma and hippocampal miR-200a-3p levels.

Hippocampal *Keap1* mRNA levels were significantly higher in T2DM mice than in control mice (Fig. 4A). They exhibited a tendency to decrease with light-intensity exercise intervention (Fig. 4A; effects of diabetes: *F*_(1, 33)_ = 5.12, *p* = 0.0304, effects of exercise: *F*_(1, 33)_ = 4.03, *p* = 0.0529, interaction: *F*_(1, 33)_ = 1.12, *p* = 0.2986). Sedentary ob/ob mice exhibited significantly higher hippocampal *Keap1* mRNA levels than exercised C57BL/6 mice (Fig. 4A). A trend of negative correlation was observed between the levels of miR-200a-3p and *Keap1* mRNA levels in the hippocampus of ob/ob mice (Fig. 4E). Hippocampal *Nrf2* mRNA levels remained unaffected by T2DM or light-intensity exercise intervention (Fig. 4B; effects of diabetes: *F*_(1, 33)_ = 0.14, *p* = 0.7099, effects of exercise: *F*_(1, 33)_ = 2.37, *p* = 0.1335, interaction: *F*_(1, 33)_ = 0.0003, *p* = 0.9866). However, only sedentary ob/ob mice exhibited significantly lowered *Hsp90aa1* mRNA levels in the hippocampus compared to sedentary C57BL/6 mice (Fig. 4C; effects of diabetes: *F*_(1, 33)_ = 7.45, *p* = 0.0101, effects of exercise: *F*_(1, 33)_ = 0.48, *p* = 0.4957, interaction: *F*_(1, 33)_ = 2.27, *p* = 0.1413). Although hippocampal *Pten* mRNA levels were significantly downregulated with light-intensity exercise intervention (Fig. 4D; effects of diabetes: *F*_(1, 33)_ = 0.04, *p* = 0.8358, effects of exercise: *F*_(1, 33)_ = 9.46, *p* = 0.0042, interaction: *F*_(1, 33)_ = 1.83, *p* = 0.1853), no correlation was found between these levels and hippocampal miR-200a-3p levels in ob/ob mice (Fig. 4F).

**Fig. 4.**
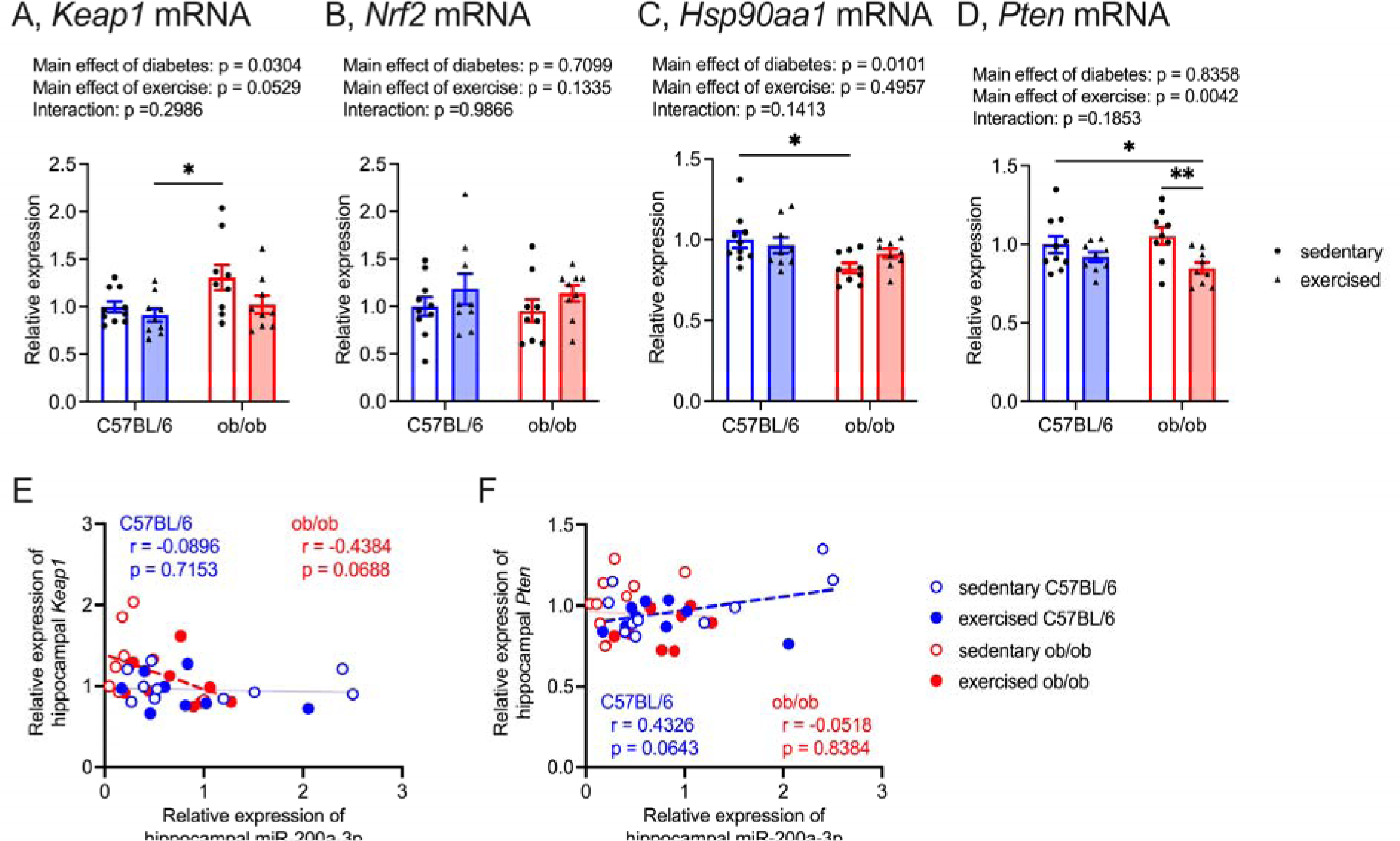
Effect of light-intensity exercise on mRNA levels of *Keap1* (A), *Nrf2* (B), *Hsp90aa1* (C), *Pten* (D) in the hippocampus. Sedentary C57BL/6 group was normalized as 100%. Data are expressed as mean ± SEM, n = 10 mice for sedentary C57BL/6, and n = 9 mice for exercised C57BL/6, sedentary ob/ob, and exercised ob/ob. \**p* < 0.05, \*\**p* < 0.01, \*\*\**p* < 0.001. (E) The correlations between miR-200a-3p levels and *Keap1* mRNA levels in hippocampus. The red dashed line in the scatter diagram indicates a trend of correlation in ob/ob mice. (F) The correlations between miR-200a-3p levels and *Keap1* mRNA levels in hippocampus. The blue dashed line in the scatter diagram indicates a trend of correlation in C57BL/6 mice.

### 3.5. The effects of intraperitoneal injection of miR-200a-3p mimic on physiological and biochemical variables

Both miRNA mimic NC-treated and miR-200a-3p mimic-treated ob/ob mice exhibited increased body weight, fat-to-body weight ratio, elevated blood glucose levels, and higher HbA_1C_ levels compared to miRNA mimic NC-treated and miR-200a-3p mimic-treated C57BL/6 mice (Table S3). No difference in these parameters was observed between miRNA mimic NC-treated and miR-200a-3p mimic-treated ob/ob mice (body weight, effects of diabetes: *F*_(1, 28)_ = 578.2, *p* < 0.0001, effects of miR-200a-3p: *F*_(1, 28)_ = 0.25, *p* = 0.6231, interaction: *F*_(1, 28)_ = 0.002, *p* = 0.9698; fat-to-body weight ratio, effects of diabetes: *F*_(1, 28)_ = 1742.2, *p* < 0.0001, effects of miR-200a-3p: *F*_(1, 28)_ = 1.51, *p* = 0.2293, interaction: *F*_(1, 28)_ = 0.35, *p* = 0.5570; blood glucose levels, effects of diabetes: *F*_(1, 28)_ = 35.0, *p* < 0.0001, effects of miR-200a-3p: *F*_(1, 28)_ = 2.85, *p* = 0.1023, interaction: *F*_(1, 28)_ = 2.74, *p* = 0.1091; HbA_1C_ levels, effects of diabetes: *F*_(1, 28)_ = 167.4, *p* < 0.0001, effects of miR-200a-3p: *F*_(1, 28)_ = 0.06, *p* = 0.8101, interaction: *F*_(1, 28)_ = 0.13, *p* = 0.7187).

### 3.6. The changes in memory function with intraperitoneal injection of miR-200a-3p mimic

On 3^rd^ day of learning session, ob/ob mice treated with miRNA mimic NC exhibited notably prolonged escape latency compared to both C57BL/6 mice treated with miRNA mimic NC and miR-200a-3p mimic; conversely, ob/ob mice treated with miR-200a-3p mimic did not exhibit this prolonged escape latency (Fig. 5A; effects of day: *F*_(3, 372)_ = 3.88, *p* = 0.0094, effects of diabetes: *F*_(1, 124)_ = 118.5, *p* < 0.0001, effects of miR-200a-3p: *F*_(1, 124)_ = 4.50, *p* = 0.0360, interaction [day × diabetes]: *F*_(3, 372)_ = 4.72, *p* = 0.0030, interaction [day × miR-200a-3p]: *F*_(3, 372)_ = 0.25, *p* = 0.8582, interaction [diabetes × miR-200a-3p]: *F*_(1, 124)_ = 0.03, *p* = 0.8620, interaction [day × diabetes × miR-200a-3p]: *F*_(3, 372)_ = 0.05, *p* = 0.9862). Swim distance and swimming speed were influenced by time and T2DM, yet treatment with miR-200a-3p mimic did not impact these parameters (Fig. 5B and C; swim distance, effects of day: *F*_(3, 372)_ = 15.42, *p* < 0.0001, effects of diabetes: *F*_(1, 124)_ = 16.76, *p* < 0.0001, effects of miR-200a-3p: *F*_(1, 124)_ = 0.64, *p* = 0.4262, interaction [day × diabetes]: *F*_(3, 372)_ = 0.16, *p* = 0.9225, interaction [day × miR-200a-3p]: *F*_(3, 372)_ = 0.30, *p* = 0.8228, interaction [diabetes × miR-200a-3p]: *F*_(1, 124)_ = 2.81, *p* = 0.0962, interaction [day × diabetes × miR-200a-3p]: *F*_(3, 372)_ = 0.17, *p* = 0.9195; swimming speed, effects of day: *F*_(3, 372)_ = 10.31, *p* < 0.0001, effects of diabetes: *F*_(1, 124)_ = 175.8, *p* < 0.0001, effects of miR-200a-3p: *F*_(1, 124)_ = 0.13, *p* = 0.7176, interaction [day × diabetes]: *F*_(3, 372)_ = 5.88, *p* = 0.0006, interaction [day × miR-200a-3p]: *F*_(3, 372)_ = 0.99, *p* = 0.3993, interaction [diabetes × miR-200a-3p]: *F*_(1, 124)_ = 1.47, *p* = 0.2272, interaction [day × diabetes × miR-200a-3p]: *F*_(3, 372)_ = 0.17, *p* = 0.9144). Furthermore, akin to light-intensity exercise intervention, the daily intraperitoneal injection of miR-200a-3p mimic increased the times spent in the platform area during the probe test for ob/ob mice (Fig. 5D). Conversely, ob/ob mice treated with miRNA mimic NC exhibited significantly shorter times spent in the platform area compared to C57BL/6 mice treated with miR-200a-3p mimic (Fig. 5D; effects of diabetes: *F*_(1, 28)_ = 0.07, *p* = 0.7989, effects of miR-200a-3p: *F*_(1, 28)_ = 10.07, *p* = 0.0036, interaction: *F*_(1, 28)_ = 4.31, *p* = 0.0473).

**Fig. 5.**
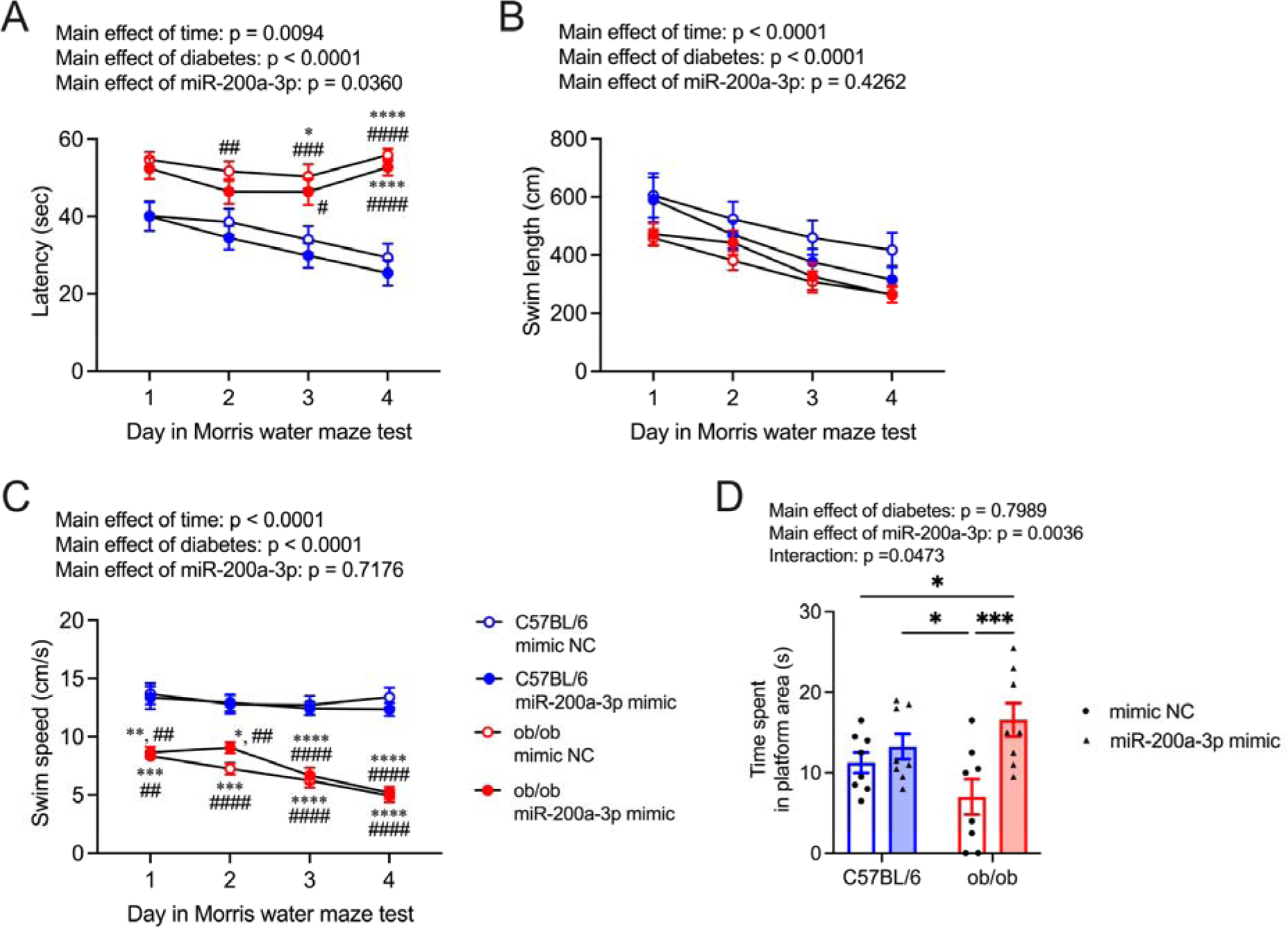
Effect of daily i.p. injection of mmu-miR-200a-3p mimic on memory function. Escape latency (A), swim length (B), and speed (C) during the learning session in mice (mean ± SEM). Blue circles: C57BL/6 mice, red circles: ob/ob mice, mimic NC: mice treated miRNA mimic negative control, miR-200a-3p mimic, mice treated mmu-miR-200a-3p mimic. \**p* < 0.05, \*\**p* < 0.01, \*\*\*\**p* < 0.0001 vs C57BL/6 mimic NC, ^#^*p* < 0.05, ^##^*p* < 0.01, ^###^*p* < 0.001, ^####^*p* < 0.0001 vs C57BL/6 miR-200a-3p mimic. (D) Effect of exercise on the probe trial, showing the crossing times where the platform had been placed. Blue bars: C57BL/6 mice, red bars: ob/ob mice, circles: mice treated miRNA mimic negative control, and triangles: mice treated mmu-miR-200a-3p mimic. \**p* < 0.05, \*\*\**p* < 0.001. Data are expressed as mean ± SEM, n = 8 mice for each group.

### 3.7. The changes in mRNA levels in the hippocampus with intraperitoneal injection of miR-200a-3p mimic

Hippocampal mRNA levels of *Mct1* and *Mct4* remained unaffected by T2DM or miR-200a-3p mimic treatment (Fig. 6A and C; *Mct1*, effects of diabetes: *F*_(1, 28)_ = 0.003, *p* = 0.9576, effects of miR-200a-3p: *F*_(1, 28)_ = 3.83, *p* = 0.0603, interaction: *F*_(1, 28)_ = 0.34, *p* = 0.5636; *Mct4*, effects of diabetes: *F*_(1, 28)_ = 1.15, *p* = 0.2931, effects of miR-200a-3p: *F*_(1, 28)_ = 0.77, *p* = 0.3876, interaction: *F*_(1, 28)_ = 3.40, *p* = 0.0757). However, the daily intraperitoneal injection of miR-200a-3p mimic notably improved *Mct2* mRNA levels in the hippocampus of ob/ob mice (Fig. 6B; effects of diabetes: *F*_(1, 28)_ = 15.8, *p* = 0.0004, effects of miR-200a-3p: *F*_(1, 28)_ = 1.81, *p* = 0.1896, interaction: *F*_(1, 28)_ = 0.21, *p* = 0.6487). Furthermore, only ob/ob mice treated with miRNA mimic NC exhibited downregulated levels of *Mct2* mRNA compared to C57BL/6 mice (Fig. 6B). Hippocampal *Hcar1* mRNA levels were significantly higher in T2DM mice than in control mice (Fig. 6D; effects of diabetes: *F*_(1, 28)_ = 33.9, *p* < 0.0001, effects of miR-200a-3p: *F*_(1, 28)_ = 2.26, *p* = 0.1438, interaction: *F*_(1, 28)_ = 0.01, *p* = 0.9158). In contrast, both miRNA mimic NC- and miR-200a-3p mimic-treated ob/ob mice exhibited significantly lower *Bdnf* mRNA levels in the hippocampus compared to C57BL/6 mice treated miRNA mimic NC (Fig. 6E; effects of diabetes: *F*_(1, 28)_ = 16.5, *p* = 0.0004, effects of miR-200a-3p: *F*_(1, 28)_ = 0.09, *p* = 0.7656, interaction: *F*_(1, 28)_ = 1.15, *p* = 0.2938). Hippocampal *Trkb* and *Creb1* mRNA levels remained unchanged by T2DM or miR-200a-3p mimic treatment (Fig. 6F and G; *Trkb*, effects of diabetes: *F*_(1, 28)_ = 0.02, *p* = 0.8923, effects of miR-200a-3p: *F*_(1, 28)_ = 0.04, *p* = 0.8458, interaction: *F*_(1, 28)_ = 1.44, *p* = 0.2408; *Creb1*, effects of diabetes: *F*_(1, 28)_ = 2.35, *p* = 0.1362, effects of miR-200a-3p: *F*_(1, 28)_ = 1.20, *p* = 0.2818, interaction: *F*_(1, 28)_ = 0.09, *p* = 0.7732). Ob/ob mice treated miRNA mimic NC solely exhibited upregulated levels of *Keap1* mRNA compared to all other groups (Fig. 7A; effects of diabetes: *F*_(1, 28)_ = 11.5, *p* = 0.0021, effects of miR-200a-3p: *F*_(1, 28)_ = 3.91, *p* = 0.0578, interaction: *F*_(1, 28)_ = 7.05, *p* = 0.0129), and downregulated levels of *Hsp90aa1* mRNA compared to C57BL/6 mice treated miRNA mimic NC (Fig. 7C; effects of diabetes: *F*_(1, 28)_ = 8.61 *p* = 0.0066, effects of miR-200a-3p: *F*_(1, 28)_ = 0.07, *p* = 0.7929, interaction: *F*_(1, 28)_ = 1.59, *p* = 0.2183). Hippocampal *Nrf2* and *Pten* mRNA levels remained unaltered by T2DM or miR-200a-3p mimic treatment (Fig. 7B and D; *Nrf2*, effects of diabetes: *F*_(1, 28)_ = 0.35 *p* = 0.5589, effects of miR-200a-3p: *F*_(1, 28)_ = 1.28, *p* = 0.2670, interaction: *F*_(1, 28)_ = 3.33, *p* = 0.0787; *Pten*, effects of diabetes: *F*_(1, 28)_ = 0.15 *p* = 0.6981, effects of miR-200a-3p: *F*_(1, 28)_ = 0.90, *p* = 0.3506, interaction: *F*_(1, 28)_ = 2.97, *p* = 0.0959).

**Fig. 6.**
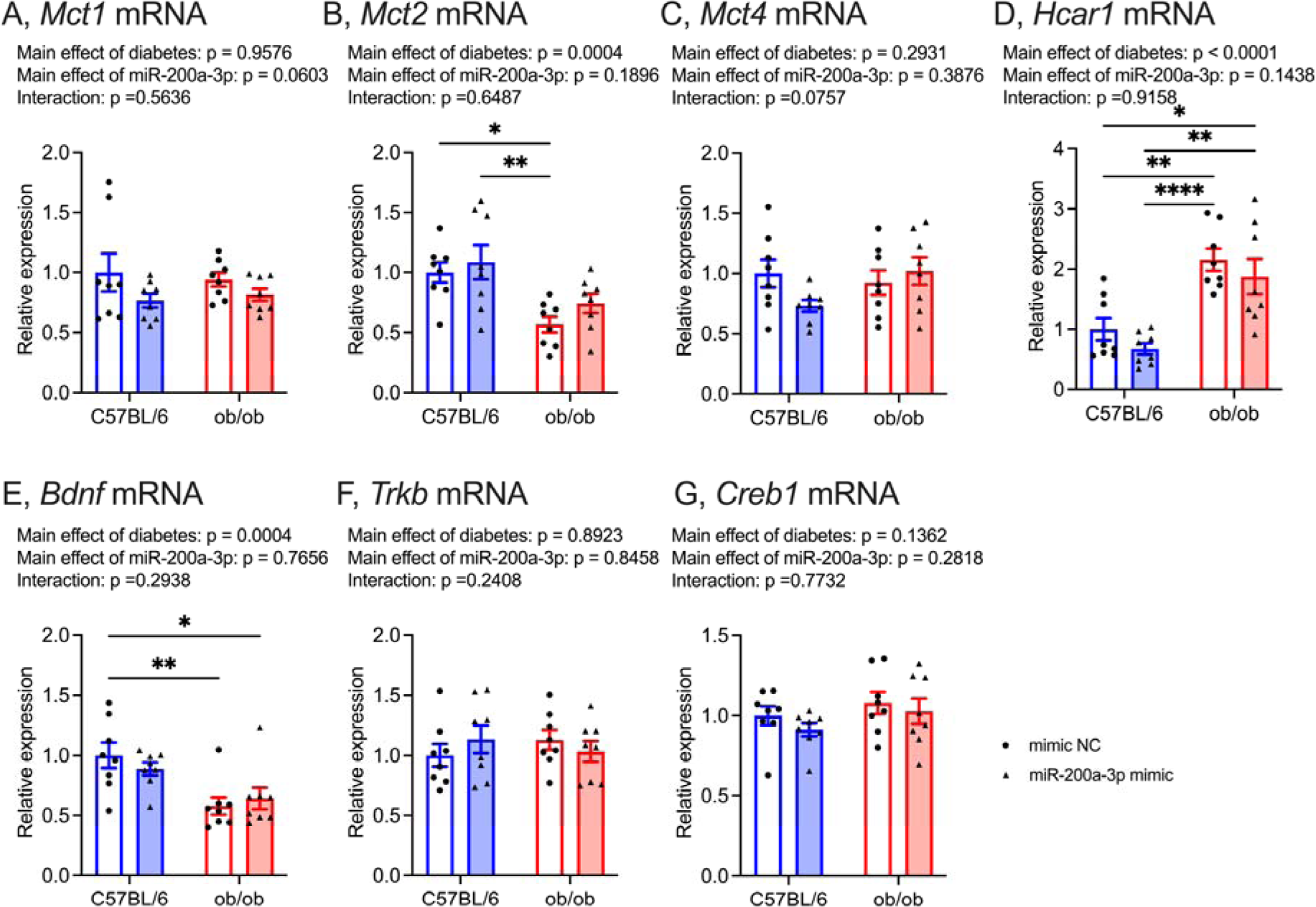
Effect of daily i.p. injection of mmu-miR-200a-3p mimic on mRNA levels of *Mct1* (A), *Mct2* (B), *Mct4* (C), *Hcar1* (D), *Bdnf* (E), *Trkb* (F), *Creb1* (G) in the hippocampus. C57BL/6 mimic NC group was normalized as 100%, mimic NC: mice treated miRNA mimic negative control, miR-200a-3p mimic, mice treated mmu-miR-200a-3p mimic. Data are expressed as mean ± SEM, n = 8 mice for each group. \**p* < 0.05, \*\**p* < 0.01, \*\*\**p* < 0.001.

**Fig. 7.**
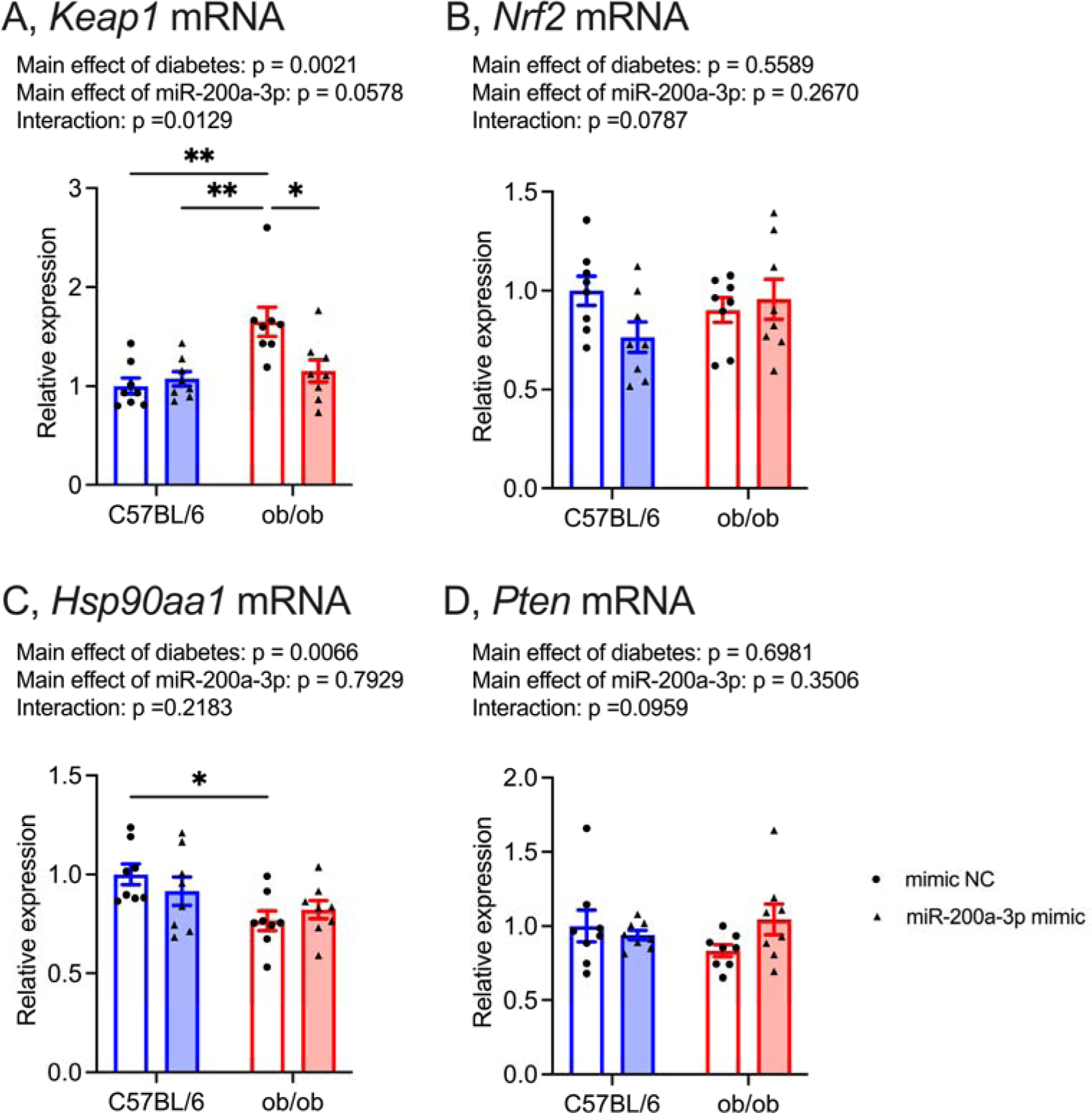
Effect of daily i.p. injection of mmu-miR-200a-3p mimic on mRNA levels of *Keap1* (A), *Nrf2* (B), *Hsp90aa1* (C), *Pten* (D) in the hippocampus. C57BL/6 mimic NC group was normalized as 100%, mimic NC: mice treated miRNA mimic negative control, miR-200a-3p mimic, mice treated mmu-miR-200a-3p mimic. Data are expressed as mean ± SEM, n = 8 mice for each group. \**p* < 0.05, \*\**p* < 0.01.

## 4. Discussion

In the current study, we investigated the involvement of muscle-derived exosomal miR-200a-3p in the improvements of memory dysfunction in T2DM mice undergoing light-intensity exercise intervention. Our findings indicated that light-intensity exercise intervention restores gastrocnemius muscle-derived exosomal miR-200a-3p levels, accompanied by improvements in spatial memory function and hippocampal *Mct2* mRNA levels in ob/ob mice. Additionally, the exercise intervention downregulated *Keap1* mRNA levels and upregulated *Hsp90aa1* mRNA levels in the hippocampus of ob/ob mice. Furthermore, the daily intraperitoneal injection of miR-200a-3p mimic in ob/ob mice not only alleviated memory dysfunction but also mimicked the effects on the hippocampal RNA profiles observed with light-intensity exercise intervention.

Consistent with previous reports [11,23], the current study reaffirms the positive impact of light-intensity exercise intervention on memory dysfunction and hippocampal *Mct2* mRNA levels in T2DM mice (Fig. 1 and 2B). The mRNA levels of Mct1 and Mct4 in the hippocampus, responsible for releasing lactate into the extracellular fluid from astrocytes, remained unaffected by both T2DM and the current light-intensity exercise regimen (Fig. 2A and C). Given that MCT2 primarily facilitates the uptake of lactate into neurons [15,45,46]. The lactate transport through MCT2 is crucial for maintaining hippocampal function [16,47,48]; restoring hippocampal *Mct2* levels might partially contribute to exercise-induced improvement of memory dysfunction in T2DM mice. Interestingly, mRNA levels of *Hcar1*, a specific receptor for lactate, were upregulated with T2DM (Fig. 2D). HCAR1, also known as GPR81, could play roles in enhancing neuroplasticity and in exercise-induced hippocampal adaptations [49–51]. Despite T2DM mice exhibiting hippocampus-based memory dysfunction, their *Hcar1* mRNA levels were higher than those of control mice; notably, hippocampal *Hcar1* mRNA levels were unchanged with light-intensity exercise intervention. While further investigations are warranted to explore the sensitivity of HCAR1 in T2DM, our findings suggest the importance of MCT2 in the exercise-induced improvement of memory dysfunction in T2DM.

Aligning with the improvement of memory dysfunction and hippocampal *Mct2* mRNA levels in T2DM mice, there was an improvement of gastrocnemius muscle-derived exosomal miR-200a-3p, exosomal miR-200a-3p in plasma, and hippocampal miR-200a-3p (Fig. 3A-C). The current study, in accordance with previous report [11], affirms the enhancement of miR-200a-3p levels in the hippocampus of T2DM mice with the light-intensity exercise regimen. Furthermore, the current results unveiled a significant positive relationship between exosomal miR-200a-3p secreted from gastrocnemius muscle and those in plasma (Fig. 3D). While exosomal miR-200a-3p levels in plasma exhibited nonsignificant correlation with hippocampal miR-200a-3p levels (Fig. 3E), the daily intraperitoneal injection of mmu-miR-200a-3p mimic improved memory dysfunction and hippocampal *Mct2* mRNA levels in T2DM mice (Fig. 5 and 6B); thus, there is a possibility that transporting peripheral miR-200a-3p into hippocampus contributes to maintaining hippocampal health in T2DM. It would be essential to explore exosome secretion from other types of muscles or organs; our current findings imply a novel interaction between the gastrocnemius muscle and the hippocampus with light-intensity exercise in T2DM.

The current exercise regimen in T2DM mice led to the downregulation of hippocampal *Keap1* and *Pten* mRNA levels (Fig. 4A, D). Although both *Keap1* and *Pten* mRNA expressions are regulated by miR-200a-3p [29,30], hippocampal miR-200a-3p expressions in T2DM mice exhibited a trend of negative correlation with *Keap1* mRNA levels, but not with *Pten* mRNA levels (Fig. 4F, G). Furthermore, mmu-miR-200a-3p mimic downregulated hippocampal *Keap1* mRNA levels, but not *Pten* mRNA levels in T2DM mice (Fig. 7A, D). Our findings suggest that the improvement of miR-200a-3p expressions in hippocampus induced by light-intensity exercise would contribute to downregulating hippocampal *Keap1* mRNA levels in T2DM. KEAP1 inhibits the expression of NRF2 and HSP90 [31,32]. The current exercise intervention improved *Hsp90aa1* mRNA levels, but not *Nrf2* mRNA levels, in the T2DM hippocampus (Fig. 4B, C). HSP90 could contribute to enhancing hippocampal neuroplasticity [52,53]. Patients with Alzheimer’s disease (AD) exhibit lower levels of HSP90 in their hippocampus compared to subjects without AD [54], and HSP90 may be related to the association between T2DM and AD [55]. The treatment of mmu-miR-200a-3p mimic led to changes in hippocampal *Hsp90aa1* mRNA levels in T2DM mice, similar to the effects of light-intensity exercise intervention (Fig. 7B); thus, we suggest that KEAP1/HSP90 signaling modulated by miR-200a-3p may contribute to improving hippocampal dysregulation in T2DM. Although the profiles of HSPs relate to the changes in expressions of MCTs [38,39], the direct relationship between HSP90 and MCT2 is unclear. Further mechanism-based studies are needed in this regard.

Consistent with previous reports [56,57], hippocampal *Bdnf* mRNA levels were downregulated by T2DM (Fig. 2E and 6E). BDNF expression in the brain is known to be regulated by PTEN [37]; however, the alterations in *Pten* mRNA associated with T2DM or exercise were inconsistent with the alterations of *Bdnf* mRNA in hippocampus (Fig. 2E, 4E, 6E, 7E). Thus, downregulated expression of BDNF in T2DM might not be directly linked to PTEN. BDNF modulates MCT2 expressions [40,41], but hippocampal *Bdnf* mRNA levels and its downstream (*Trkb* and *Creb1* mRNA) in T2DM mice remained unchanged with the current light-intensity exercise (Fig. 2E-G) or treatment of mmu-miR-200a-3p mimic (Fig. 6E-G), implying that the light-intensity exercise restored memory function and hippocampal *Mct2* levels regardless of BDNF.

As shown in previous studies [11,58,59], elevated blood glucose and HbA_1C_ levels in T2DM mice were improved with light-intensity exercise (Table S2); however, the daily intraperitoneal injection of mmu-miR-200a-3p mimic did not alter these biochemical parameters (Table S3). Therefore, it appears that miR-200a-3p may not have a direct impact on improving glycometabolism in peripheral organs. Instead, miR-200a-3p might represent a specific therapeutic strategy for targeting the hippocampus in T2DM.

The current study has some limitations. Firstly, there is a need for further investigation into the specific mechanisms and interactions underlying how the uptake of lactate through hippocampal neuronal MCT2 is linked to memory dysfunction in T2DM. Secondly, our study focused solely on the hippocampal lactate transporter, neglecting the measurement of other molecular factors implicated in T2DM-induced memory dysfunction, including angiogenesis, inflammation, oxidative stress, and insulin resistance [6,60,61]. For instance, a previous study has demonstrated a link between hippocampal insulin resistance and the downregulation of neuroplasticity and cognitive function [62]. Given that drugs enhancing insulin sensitivity and intranasal insulin injection have been shown to improve hippocampus-based memory function in T2DM [63–65], hippocampal insulin resistance would be an important target to treat cognitive dysfunction associated with T2DM. Therefore, further studies are warranted to investigate the impact of the current interventions on insulin action in the hippocampus. Thirdly, the current study exclusively assessed hippocampal spatial memory function using the Morris water maze test, thereby leaving the effects of the interventions on other forms of hippocampus-based memory function, such as novel object recognition, yet to be elucidated. Additionally, memory function and circulating biochemical parameters in mice were evaluated only after the interventions, with no assessments conducted prior to the interventions. Fourthly, our specific emphasis on the modulation of *Keap1* and *Pten* mRNA by miR-200a-3p may not capture the full spectrum of its effects, as some studies have reported adverse outcomes, such as the promotion of neuronal apoptosis by miR-200a-3p [66]. Thus, a more comprehensive exploration of these aspects is warranted in future investigations. Finally, our measurements were restricted to mRNA levels, and we lack information on protein levels in the hippocampus. Based solely on the current findings, it is impossible to completely reveal the specific pathway of the interventions’ effects. Hence, further mechanism-based studies, including genetic modulations and ex-vivo investigations, are warranted.

In conclusion, our current findings represented that light-intensity exercise intervention in T2DM mice effectively ameliorates memory dysfunction and enhances hippocampal *Mct2* mRNA levels, accompanied by the improvement of gastrocnemius muscle-derived exosomal miR-200a-3p levels. The observed downregulation of hippocampal *Keap1* mRNA and upregulation of *Hsp90aa1* mRNA in response to both light-intensity exercise and miR-200a-3p mimic treatment in T2DM mice suggest a potential involvement of the KEAP1/HSP90 signaling pathway modulated by miR-200a-3p in improving hippocampal dysregulation. These findings have the potential to advance and innovate therapeutic approaches for enhancing hippocampal health in T2DM.

## Funding

This work was supported by the Japan Society for the Promotion of Science (Grant-in-Aid for Early-Career Scientists, No. JP20K19565; JP22K17711).

### Data availability

The datasets in the current study are available from the corresponding author on reasonable request.

### CRediT authorship contribution statement

**Takeru Shima**: Conceptualization, Methodology, Resources, Investigation, Writing– original draft, Writing–review & editing, Funding acquisition; **Hayate Onishi**: Investigation, Writing–review & editing; **Chiho Terashima**: Investigation, Writing–review & editing.

## Abbreviations

BDNF: brain-derived neurotrophic factor
CREB: cAMP response element binding protein
HSP90: heat shock protein 90
KEAP1: kelch ECH-associated protein 1
MCT: monocarboxylate transporter
NRF2: nuclear factor erythroid 2 p45-related factor-2
PTEN: phosphatase and tensin homolog
T2DM: Type 2 diabetes mellitus
TrkB: tropomyosin[related kinase B

## Supplementary Tables

**Supplementary Table 1.**
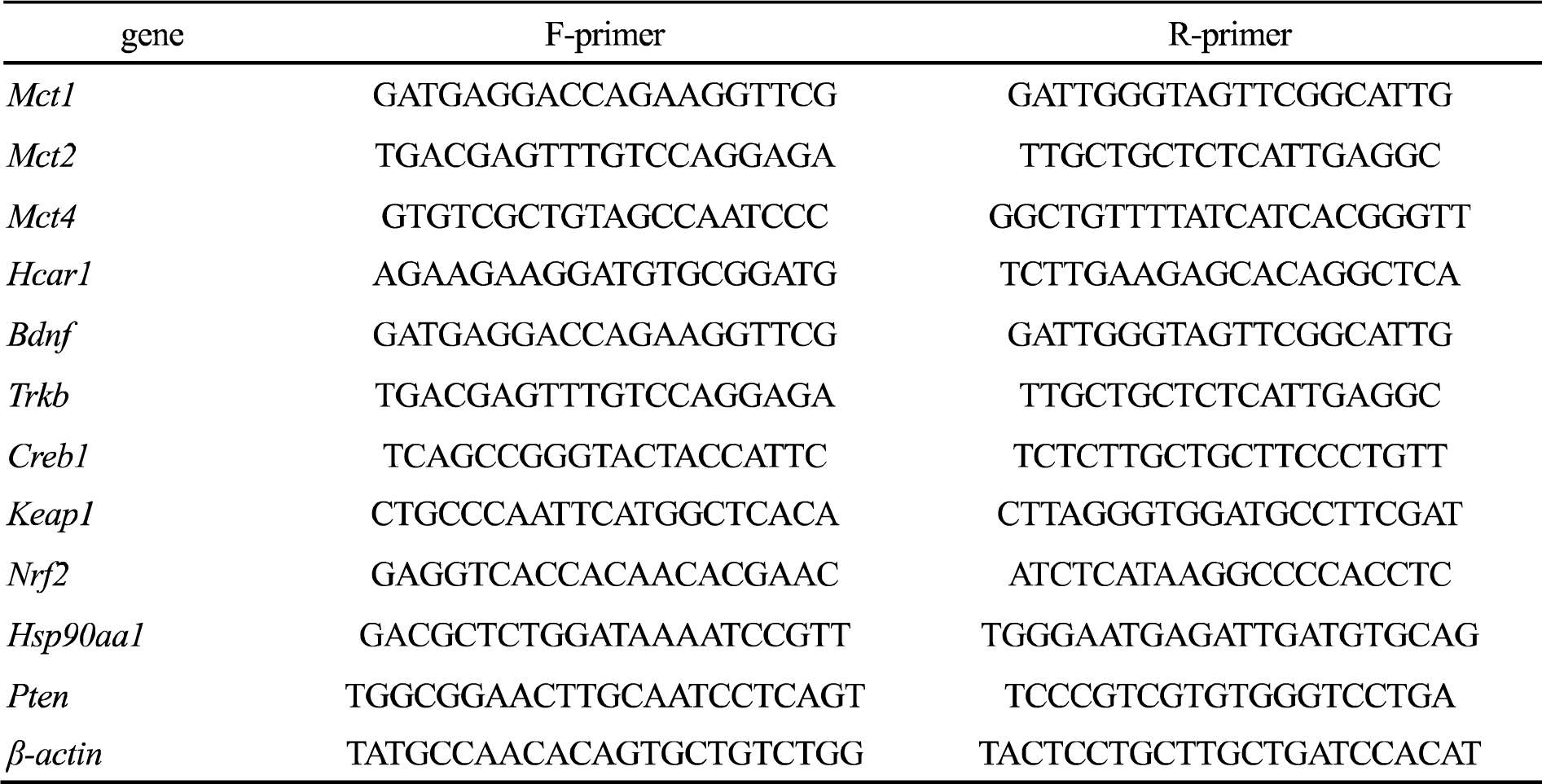
Primer sequences.

**Supplementary Table 2.** The effects of light-intensity exercise on physiological and biochemical variables.

|  | C57BL/6 |  | ob/ob |  |
| --- | --- | --- | --- | --- |
|  | sedentary | exercised | sedentary | exercised |
| Body weight (g) | 26.53 ± 0.42 | 25.19 ± 0.35 | 52.19 ± 1.19****,#### | 52.11 ± 0.62****,#### |
| Fat weight ( mg/g body weight ) | 3.82 ± 0.28 | 3.11 ± 0.17 | 55.83 ± 1.57****,#### | 52.62 ± 1.66****,#### |
| Blood glucose (mg/dL) | 145.10 ± 6.38 | 134.67 ± 6.74 | 216.11 ± 25.38**,## | 178.22 ± 12.85 |
| HbA<sub>1C</sub> (%) | 4.29 ± 0.05 | 4.27 ± 0.06 | 7.67 ± 0.26****,#### | 6.60 ± 0.17****,####,+++ |

Values are mean ± SEM. \*\* p < 0.01,\*\*\*\* p < 0.0001 vs C57BL/6-sedentary, ### p < 0.001, ##### p < 0.0001 vs C57BL/6-exercised, +++ p < 0.001 vs ob/ob-sedentary.

**Supplementary Table 3.** The effects of intraperitoneal injection of miR-200a-3p mimic on physiological and biochemical variables.

|  | C57BL/6 |  | ob/ob |  |
| --- | --- | --- | --- | --- |
|  | mimic NC | miR-200a-3p mimic | mimic NC | miR-200a-3p mimic |
| Body weight (g) | 26.43 ± 0.20 | 25.90 ± 0.13 | 53.99 ± 1.91****,#### | 53.38 ± 1.24****,#### |
| Fat weight ( mg/g body weight ) | 3.08 ± 0.25 | 2.25 ± 0.19 | 58.24 ± 2.24****,#### | 55.86 ± 1.29****,#### |
| Blood glucose (mg/dL) | 124.25 ± 9.04 | 123.50 ± 6.06 | 290.38 ± 34.71****,#### | 217.00 ± 24.55*,# |
| HbA<sub>1C</sub> (%) | 4.28 ± 0.10 | 4.24 ± 0.07 | 8.16 ± 0.42****,#### | 8.35 ± 0.44****,#### |

Values are mean ± SEM. \* p < 0.05, \*\*\*\* p < 0.0001 vs C57BL/6-mimic NC, # p < 0.05, #### p < 0.0001 vs C57BL/6-miR-200a-3p mimic.

